# Computational design of developable therapeutic antibodies: efficient traversal of binder landscapes and rescue of escape mutations

**DOI:** 10.1101/2024.10.03.616038

**Authors:** Frédéric A. Dreyer, Constantin Schneider, Aleksandr Kovaltsuk, Daniel Cutting, Matthew J. Byrne, Daniel A. Nissley, Newton Wahome, Henry Kenlay, Claire Marks, David Errington, Richard J. Gildea, David Damerell, Pedro Tizei, Wilawan Bunjobpol, John F. Darby, Ieva Drulyte, Daniel L. Hurdiss, Sachin Surade, Douglas E. V. Pires, Charlotte M. Deane

## Abstract

Developing therapeutic antibodies is a challenging endeavour, often requiring large-scale screening to produce initial binders, that still often require optimisation for developability. We present a computational pipeline for the discovery and design of therapeutic antibody candidates, which incorporates physics- and AI-based methods for the generation, assessment, and validation of developable candidate antibodies against diverse epitopes, via efficient few-shot experimental screens. We demonstrate that these orthogonal methods can lead to promising designs. We evaluated our approach by experimentally testing a small number of candidates against multiple SARS-CoV-2 variants in three different tasks: (i) traversing sequence landscapes of binders, we identify highly sequence dissimilar antibodies that retain binding to the Wuhan strain, (ii) rescuing binding from escape mutations, we show up to 54% of designs gain binding affinity to a new subvariant and (iii) improving developability characteristics of antibodies while retaining binding properties. These results together demonstrate an end-to-end antibody design pipeline with applicability across a wide range of antibody design tasks. We experimentally characterised binding against different antigen targets, developability profiles, and cryo-EM structures of designed antibodies. Our work demonstrates how combined AI and physics computational methods improve productivity and viability of antibody designs.

## 1 Introduction

Antibodies are an important and growing class of therapeutics [1], representing the largest and most successful class of biotherapeutics, with a market size estimated to reach $450B by 2028 [2]. To date, more than 100 therapies have been approved by the FDA against a wide range of disease indications [3], with particular success in oncology and immuno-oncology, and with revenues projected to surpass that of small molecule therapies in the next few years [4].

Modern antibody therapeutic discovery and design processes are heavily supplemented and accelerated by computational and machine learning (ML) driven methods [5]. These approaches include the *in silico* prediction of epitope and paratope regions [6, 7], various developability and affinity characteristics [8–10] as well as generative approaches to the development of novel candidate antibodies [11–13].

Despite recent advances in computational approaches [14–17], the majority of successful antibody drug discovery campaigns rely heavily on extensive experimental screening, initially to identify binders and in subsequent rounds to further optimise specificity, affinity, humanness, stability, and other characteristics needed for downstream development [1]. This procedure of optimising binding affinity and developability characteristics separately is typical of most discovery pipelines, which tends to drive up overall costs and prolong discovery time [18].

We propose a design pipeline that combines (i) *in silico* biophysical property assessment, (ii) machine learning-based antibody design approaches and (iii) sample-efficient experimental validation to enable the design of high affinity and developable therapeutic antibodies from starting point antibody candidates. We use our pipeline to design antibodies against multiple variants of the SARS-CoV-2 receptor binding domain (RBD), we show that our methodology is able to discover binders from naturally occurring repertoires, improve developability metrics of existing binders and rescue antibodies from escape mutations on the target antigen.

In this pilot study, we experimentally validated a small number of designs from three different computational design methods in a single-shot approach. Starting with known SARS-CoV-2 wild-type RBD-binding antibodies, we designed and characterised 285 novel antibodies against different strains of the virus. An overview of our computational design methods and characterisation pipeline is shown in Figure 1.

**Fig. 1.**
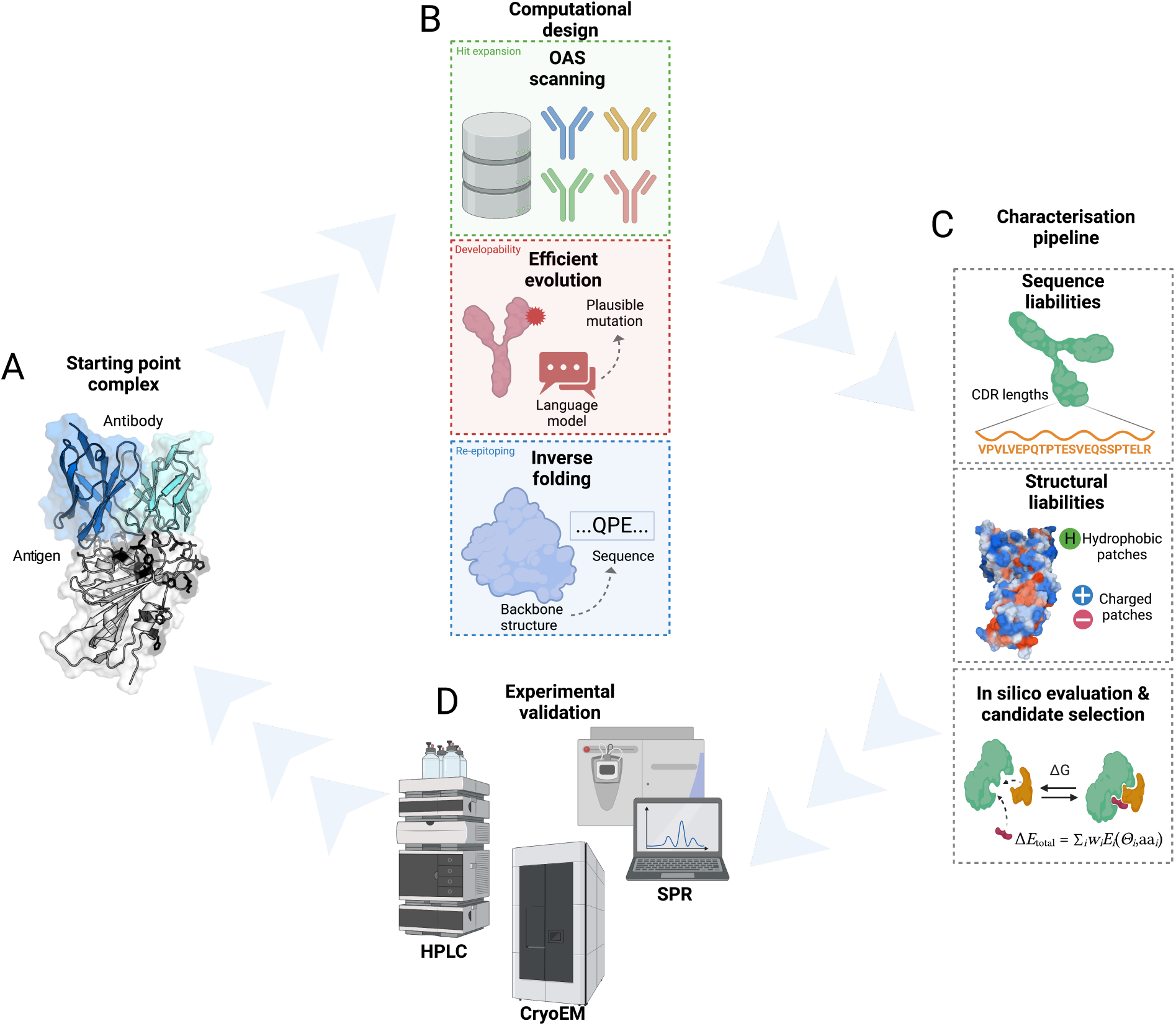
(A) An antibody-antigen complex that will be used as starting point for optimisation. (B) Overview of the computational antibody design platform, showing the three candidate generation methods. (C) The characterisation pipeline used to filter and rank candidates to select a small subset. (D) Experimental validation of selected candidates to measure properties such as binding affinity to the target, aggregation propensity and thermostability.

We demonstrate our computational pipeline is capable of (i) efficiently traversing sequence landscapes of binders, achieving high expression and hit rates through the use of effective computational screening; (ii) rescuing developability issues of existing binders, such as aggregation, through the use of language model-guided modifications; and (iii) re-epitoping [19] antibodies to mutated antigens through the use of an antibody-specific inverse folding model.

Our computational design pipeline represents a significant step towards ML-driven design of potent biotherapeutics.

## 2 Results

### 2.1 Selection of viable antibody starting points for design

We first sought a pool of template antibodies known to bind the SARS-CoV-2 RBD to serve as the basis for design. It was previously reported that antibodies binding the Class 3 epitope on RBD tend to be tolerant to mutations and thus maintain neutralisation across RBD strains [20]. We characterised a set of 192 Class 3 antibodies, of which 182 have no known structure and were selected from a larger pool of Class 3 antibodies on the basis of their broad neutralisation in a previous study [21].

To assess this pool of antibodies, we performed a measurement of binding affinity against the Wuhan SARS-CoV-2 strain, the XBB.1 strain and the XBB.1.5 strain with a single-concentration surface plasmon resonance (SPR) experiment, as described in Section 4.1.4. A summary of this study is given in Appendix A. With these results, we confirmed that eight of the ten antibodies with known structure were able to bind the original Wuhan SARS-CoV-2 strain but failed to bind, or only bound weakly, to the XBB.1.5 RBD. We selected five of these antibodies, BD55-5840, DXP-604, LY-CoV1404, REGN10987, and S2K146, which have CDR H3 loop lengths up to 15 residues, as starting points for our hit expansion and re-epitoping pipelines described in Sections 2.2 and 2.4 respectively. For our developability pipeline, in Section 2.3, we also use as an additional starting point the antibody S309, which exhibits binding to the Wuhan, XBB.1 and XBB.1.5 strains, but has poor developability characteristics, notably a propensity to aggregate and a low melting temperature. The binding affinity and developability properties of all antibodies used as starting points are given in Table 2. We resolved the structure of two of these starting point antibodies, REGN10987 and S309, using cryogenic electron microscopy as described in Section 4.3. These are shown in Figure 5 (a) and (b). Further details of the starting points used in each computational design pipeline are given in Appendix B.

### 2.2 Efficiently navigating the antibody space enables the identification of diverse binders

We took five starting point antibodies (REGN10987, BD55-5840, DXP-604, S2K146 and LY-CoV1404) with strong binding affinity against the Wuhan strain, as defined in Section 4.1.5, but no or weak binding against the XBB.1.5 strain and moderate CDR H3 loop length. These starting points were used as seeds to generate libraries of candidate antibodies from paired and unpaired sequences in the Observed Antibody Space (OAS) dataset [22, 23] as detailed in section 4.2.2. Using our characterisation pipeline, described in Section 4.2.1, to select antibodies for experimental validation, we computationally screened 11,389 candidate antibodies, of which 148 were submitted for experimental validation, 67 directly drawn from OAS and 81 designs elaborated using an inverse folding model. None of the 67 candidates drawn from OAS showed binding against the XBB.1 and XBB.1.5 strains, while 18 showed binding against the Wuhan strain in single concentration antigen SPR measurements as outlined in Section 4.1.4, with 14 strong binders and 4 medium binders, as defined in Section 4.1.5. A summary of binding and developability profile is given in Table 1, showing also the high dissimilarity of the designs to the starting point, with median edit distances from 11 to 75 depending on the starting point. About 95% of designs pass our developability criterion described in Section 4.1.6. We achieved a hit rate of 21% across the four starting points where our characterisation pipeline was able to identify binding antibodies. Of these binders, five derived from the DXP-604 and REGN10987 starting points were submitted for kinetic SPR characterisation with multiple antigen concentration, with three of these being confirmed as strong binders, as displayed in Table 3 and in Figure 2(a). Structures of the antibody-antigen complex for two designs for each starting point are displayed in Figure 2(c). We also show the full chromatograms of size exclusion chromatography in Figure 2(b). These antibodies are highly sequence-dissimilar to their respective starting point antibody, with up to 74 edits across the whole sequence. These results demonstrate that, given a starting point structure, our pipeline is able to identify sequence-distant binders with sample-efficient experimental screening.

**Fig. 2.**
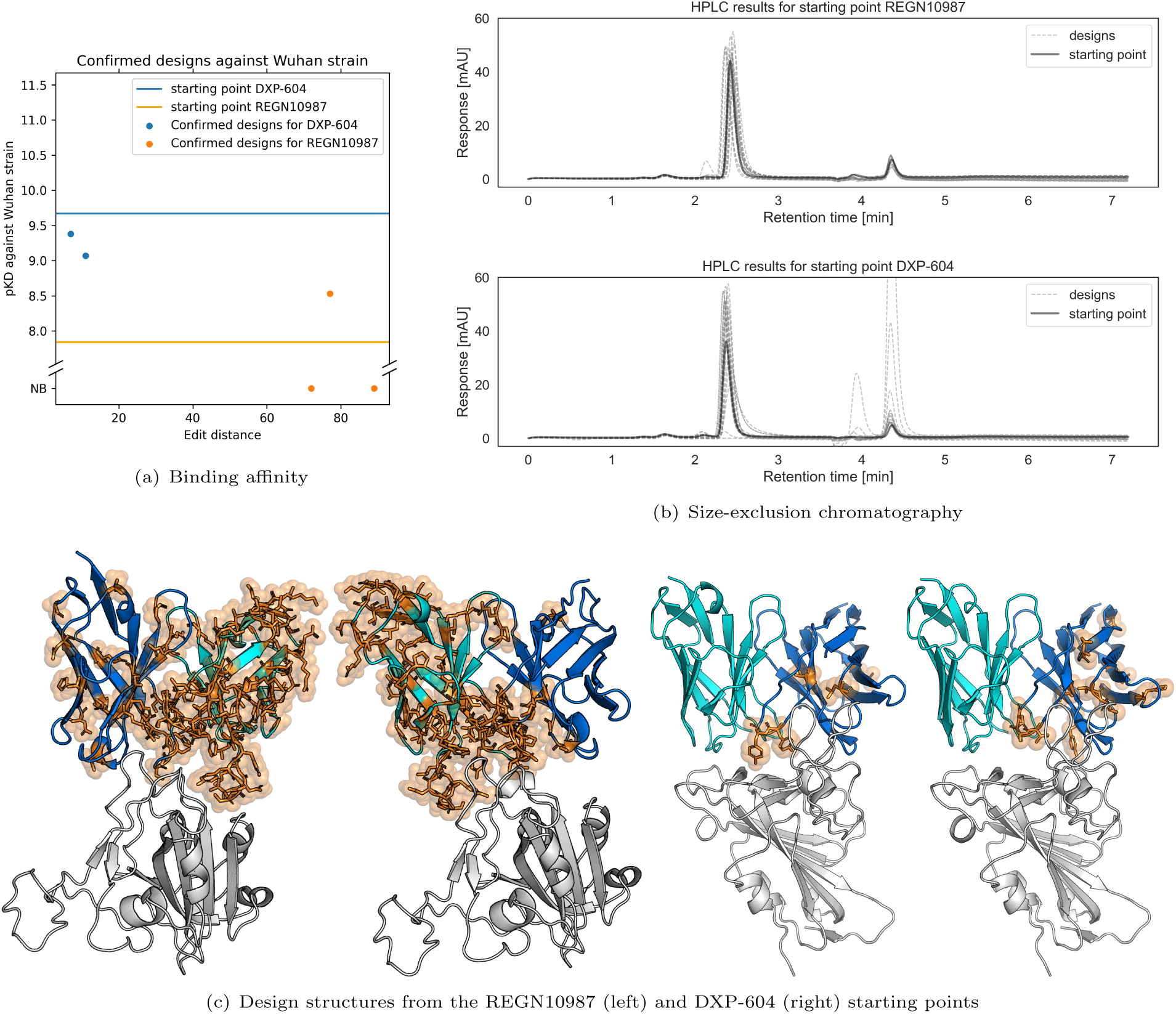
Main results of the OAS scanning design strategy. (a) Full titration SPR measurement of binding affinity to the Wuhan strain for selected designs and starting points. (b) Size-exclusion chromatography experiments for all designs and starting points. (c) Selected examples of design structures, highlighting mutations to the starting point in orange.

**Table 1.** Summary of experimental characterisation of candidate antibodies generated with our design strategies. Developable denotes designs that pass our HPLC quality threshold as detailed in Section 4.1.6. Hits are designs that show measurable binding in a single concentration antigen screen, regardless of strength of the binding interaction.

| Method | Starting point | Designs | Median edit dist. | Hits Wuhan | Hits XBB.1 | Hits XBB.1.5 | Developable |
| --- | --- | --- | --- | --- | --- | --- | --- |
| OAS scanning | BD55-5840 | 43 | 62 | 1 | 0 | 0 | 43 |
|  | DXP-604 | 26 | 11 | 22 | 1 | 1 | 23 |
|  | LY-CoV1404 | 23 | 17 | 0 | 0 | 0 | 20 |
|  | REGN10987 | 33 | 74 | 1 | 0 | 0 | 32 |
|  | S2K146 | 23 | 75 | 2 | 0 | 0 | 22 |
| Inverse folding | BD55-5840 | 13 | 11 | 1 | 0 | 0 | 3 |
|  | DXP-604 | 13 | 4 | 13 | 7 | 7 | 13 |
|  | LY-CoV1404 | 13 | 3 | 12 | 0 | 2 | 13 |
|  | REGN10987 | 13 | 10 | 2 | 0 | 0 | 12 |
|  | S2K146 | 13 | 5 | 3 | 0 | 0 | 11 |
| Efficient evolution | BD55-5840 | 12 | 7 | 12 | 0 | 0 | 12 |
|  | DXP-604 | 12 | 8 | 12 | 0 | 0 | 12 |
|  | LY-CoV1404 | 12 | 11 | 12 | 0 | 0 | 12 |
|  | REGN10987 | 12 | 9 | 2 | 0 | 0 | 10 |
|  | S2K146 | 12 | 6 | 10 | 0 | 0 | 12 |
|  | S309 | 12 | 8 | 12 | 12 | 12 | 6 |

**Table 2.**
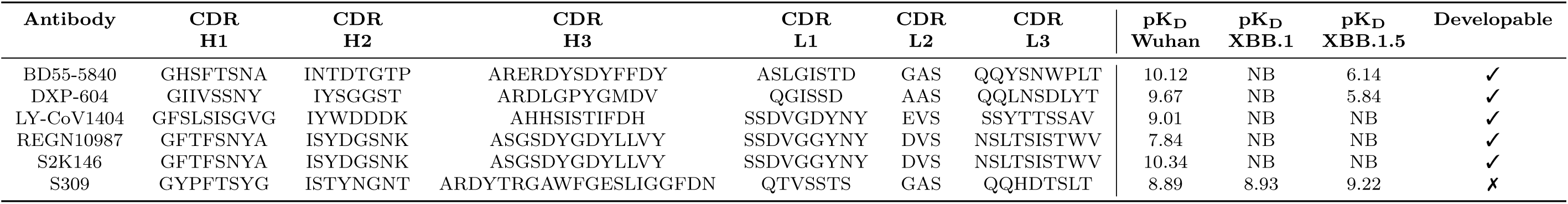
Sequence of the CDR loops, as well as kinetic characterisation with multiple antigen concentrations against three SARS-CoV-2 variants,^1^ as described in Section 4.1.4, and developability characterisation, as defined in Section 4.1.6, for each starting point antibody used in this study. ^1^ Except for S2K146 where only a single antigen concentration measurement was performed.

**Table 3.**
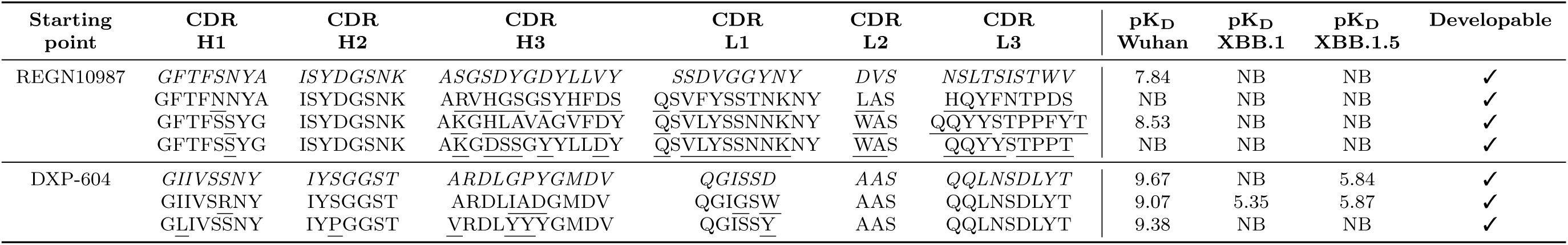
Sequence of the CDR loops, as well as kinetic characterisation with multiple antigen concentrations and developability characterisation of candidate antibodies designed with the OAS scanning strategy. The first row for each starting point contains the sequence and experimental validation of the starting point antibody itself, with mutations introduced underlined in subsequent rows.

### 2.3 Protein language model approaches enable improvement of developability properties for existing candidate antibodies

Beyond the generation of novel binders against a target of interest, improving the developability profile of existing binders while maintaining affinity to the target is also an important task in computational antibody design. Using an ensemble of six ESM language models [24] to identify sequence mutations with high likelihood [25], we generated 12,000 designs (2,000 for each of the six starting points) as described in Section 4.2.4 and characterised and ranked the designs using our characterisation pipeline.

This combination of energetics assessment with language-guided evolution allows us to improve developability properties of starting point antibodies in a single-shot *in silico* study while maintaining binding affinity with the antigen. This is in contrast with the original study [25], which required two rounds of experiments with an initial assessment of single-point mutations, and for which many antibodies lost their binding affinity to the antigen.

We submitted the 12 top ranked designs for each starting point for experimental validation, testing 72 designs in total for which a summary is given in Table 1. Of these, 57 designs maintained binding against the Wuhan strain (51 strong binders, 6 medium binders), achieving a hit rate of 79% with a median edit distance of 8. Of particular interest, the starting point antibody S309 has strong binding to the Wuhan strain as well as the XBB.1 and the XBB.1.5 strains, but has unfavorable developability characteristics, namely a propensity for aggregation [26], as measured by the size-exclusion chromatography approach described in Section 4.1.6, as well as poor thermostability. All 12 designs derived from the S309 starting point that were experimentally validated maintained binding against all three strains of the virus. We ran full-titration SPR on all 12 of the S309-derived designs and confirmed them as either strong (9) or medium (3) binders against the XBB.1.5 strain and strong binders against the Wuhan strain, as is shown in Table 4 and Figure 3(a). The antibody-antigen complex structure for a few selected designs is shown in Figure 3(c). We further ran size exclusion chromatography columns on these 12 designs. All designs showed significant improvements in aggregation and 10 of the 12 designs showed improvement in thermostability compared to the S309 starting point, as shown in the chromatograms and melting point comparisons in Figure 3(b). This demonstrates the ability of our pipeline to generate and identify candidate antibodies with favorable developability properties while maintaining existing binding affinities.

**Fig. 3.**
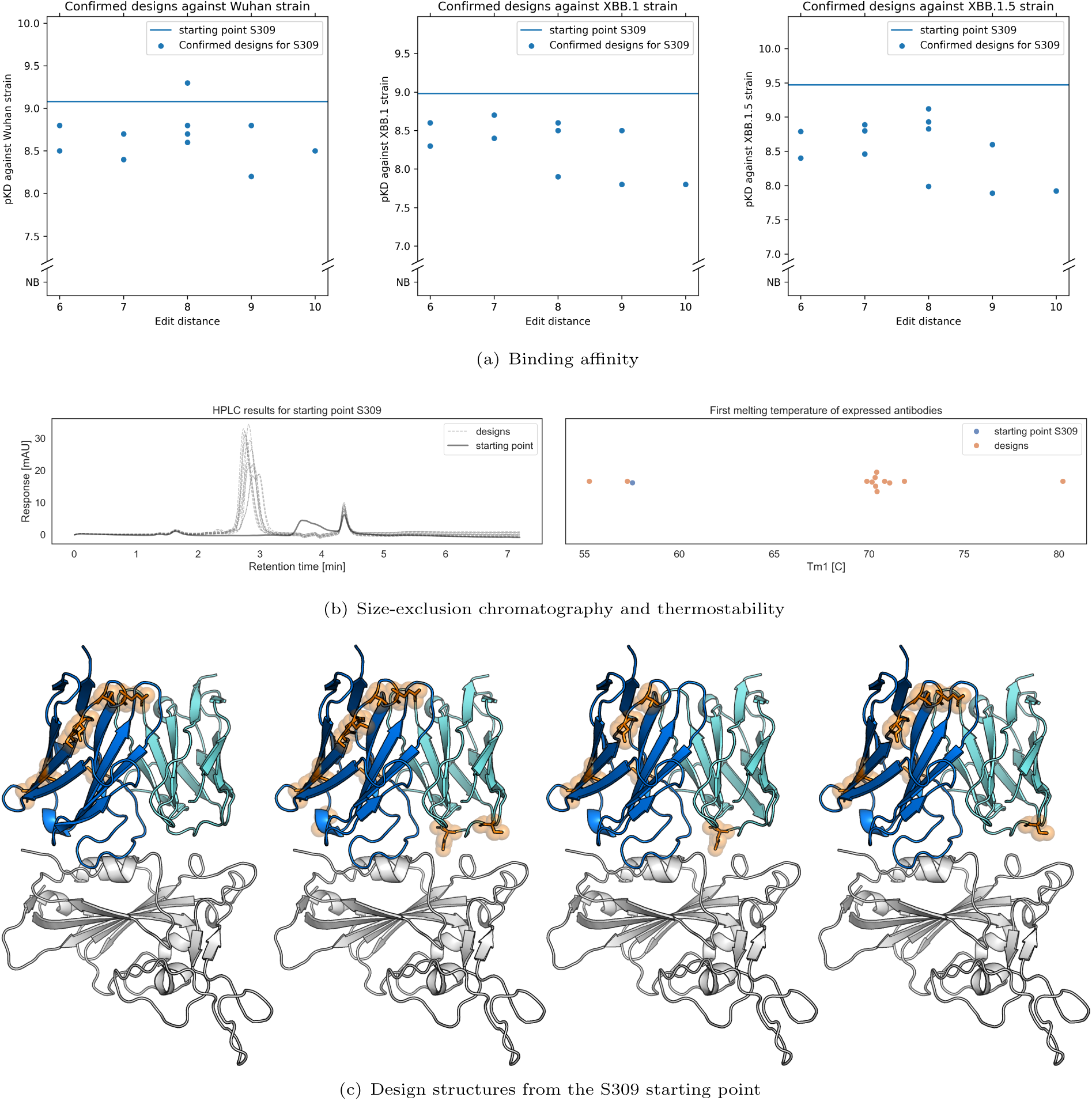
Main results of the efficient evolution design strategy. (a) Full titration SPR measurement of binding affinity to the three virus strains for selected designs and the S309 starting point. (b) Size-exclusion chromatography experiments for all designs and the starting point, as well as thermostability measurements. (c) Selected examples of design structures, highlighting mutations to the starting point in orange.

**Table 4.**
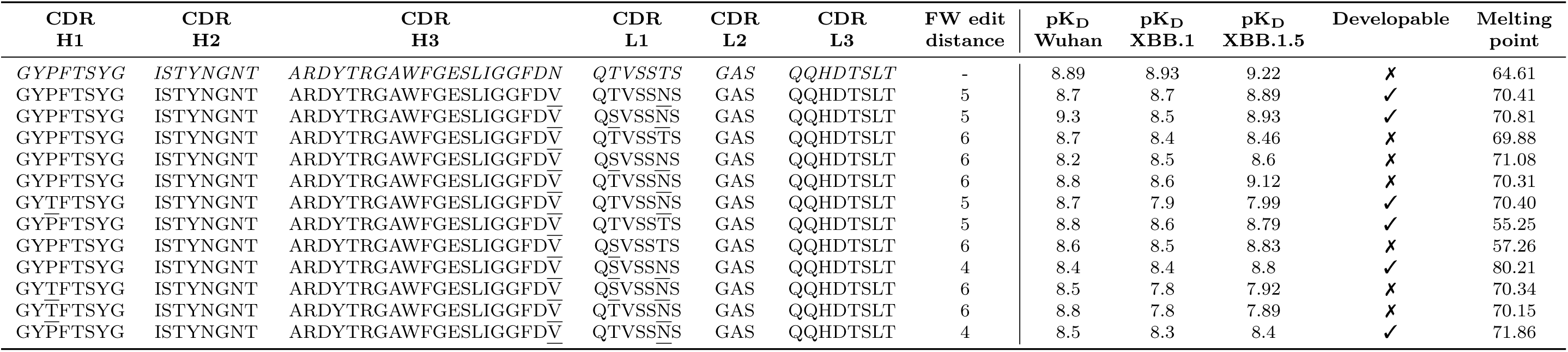
Sequence of the CDR loops, as well as kinetic characterisation with multiple antigen concentrations, thermostability and developability characterisation of candidate antibodies designed with the efficient evolution strategy. The first row contains the sequence and experimental validation of the S309 starting point antibody itself, with mutations introduced underlined in subsequent rows.

### 2.4 Inverse folding enables rescue of candidate antibodies from escape mutations

Recovering binding activity of therapeutic antibodies after escape mutations on the target antigen is a particularly important task for computational antibody design [27, 28]. We demonstrate that using an antibody-specific, antigen-aware inverse folding model and our characterisation pipeline, we can recover binding to the antigen after viral escape mutations.

To this end, we generated 25,000 designs using AbMPNN [29], as described in section 4.2.3, using five starting point antibodies (5,000 each for REGN10987, BD55-5840, DXP-604, S2K146 and LY-CoV1404). We ranked and filtered them using our characterisation pipeline. We experimentally validated the top 13 designs for each starting point through a single concentration antigen SPR measurement, of which a summary is given in Table 1. Of these 65 designs, 31 maintained binding against the Wuhan strain, with 25 strong binders and 6 medium binders, achieving a hit rate against the Wuhan strain of 48%. Amongst the designs binding to the Wuhan strain, the median edit distance is 4 with a maximum edit distance of 13.

For the starting points LY-CoV1404 and DXP-604, we demonstrate using a full-titration SPR measurement that 9 designs maintained strong binding against the Wuhan strain and gained weak (2) or medium (7) binding against XBB.1.5, while displaying favorable developability characteristics, as shown in Table 5 and in Figure 4. For the DXP-604 starting point, we achieve a hit rate of 54% against the XBB.1 and XBB.1.5 strains, while the LY-CoV1404 starting point designs have a hit rate of 15% against the XBB.1.5 strain. Furthermore, we selected one design per starting point for structural characterisation with the cryo electron microscopy methodology described in Section 4.3 to verify the binding pose of our designed antibodies against the Wuhan SARS-CoV-2 RBD. These experimentally resolved structures are shown in Figure 5 (c) and (d).

**Fig. 4.**
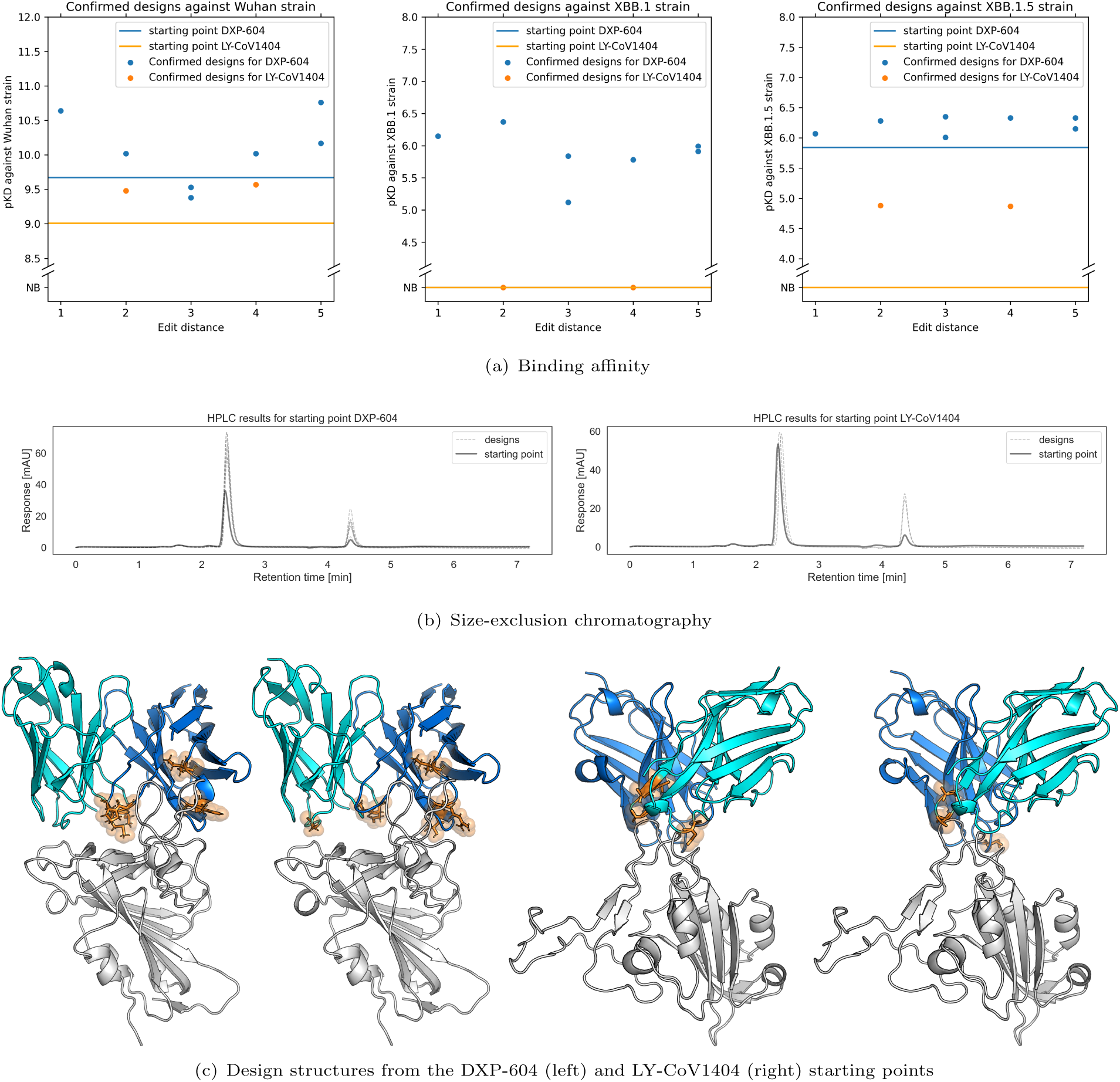
Main results of the inverse folding design strategy. (a) Full titration SPR measurement of binding affinity to the three virus strains for selected designs and starting points. (b) Size-exclusion chromatography experiments for all designs and starting points. (c) Selected examples of design structures, highlighting mutations to the starting point in orange.

**Fig. 5.**
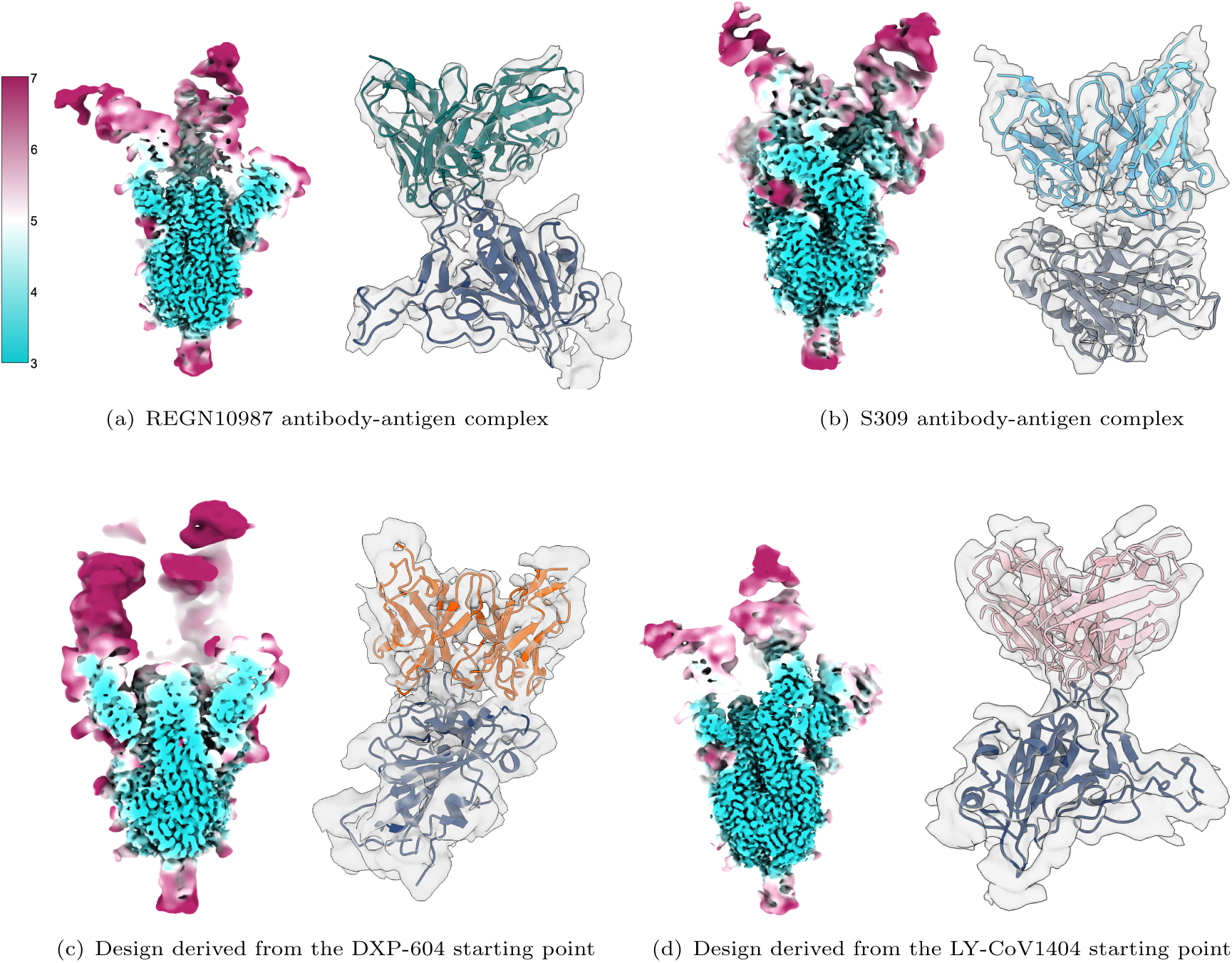
Cryo-electron microscopy epitope mapping of four selected antibodies in complex with the SARS-CoV-2 spike protein. (a) and (b) show the REGN10987 and S309 starting points, while (c) and (d) are selected designs from the inverse folding method. In each panel, the left-hand side shows central slices of 3D electron microscopy volumes coloured by local resolution in angstroms, and the right-hand side displays locally refined interfaces of the fragment antigen-binding region and receptor-binding domain (in blue).

**Table 5.**
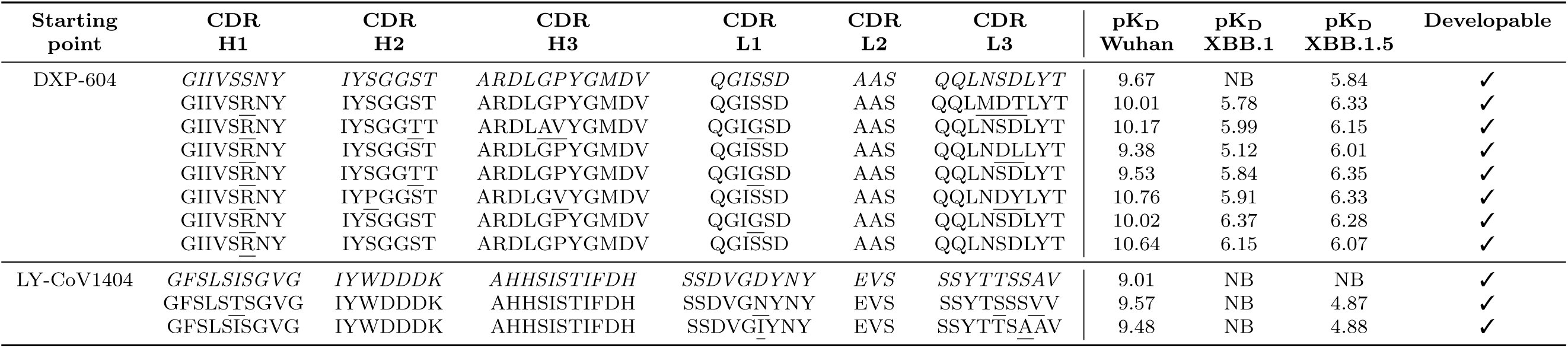
Sequence of the CDR loops, as well as kinetic characterisation with multiple antigen concentrations and developability characterisation of candidate antibodies designed with the inverse folding strategy. The first row for each starting point contains the sequence and experimental validation of the starting point antibody itself, with mutations introduced underlined in subsequent rows.

With these results, we demonstrate that our candidate antibody generation and characterisation pipeline is not only able to improve binding and developability for existing antibody binders but is also able to yield antibodies binding to previously non-bound epitopes.

## 3 Discussion

We introduce a novel pipeline for the discovery and design of potent and developable therapeutic antibody candidates. Our approach, which integrates physics-based and ML-driven antibody design and characterisation, successfully identifies effective and sequence-diverse binders with favorable developability profiles from the natural antibody repertoire, using existing antibodies as starting points. Our pipeline presents a valuable opportunity for hit elaboration during antibody design campaigns. We demonstrate that combining our characterisation pipeline with a language model for sequence-based elaboration enhances the developability profiles of candidate antibodies while maintaining binding potency in a single round of *in silico* screening. This suggests a viable strategy for the lead optimisation of antibody therapeutics. Lastly, we show that using an inverse folding model in combination with our characterisation pipeline can allow for the restoration of binding activity in candidate antibodies after antigen escape mutations, using low-sample experimental screening.

Our computational design pipeline could be combined with a Bayesian optimisation approach [30–34] to improve properties of candidate antibodies over multiple design cycles with experimental validation. Future directions for research also include incorporating recent advancements in antibody-antigen structure prediction using modern ML techniques [35, 36] to facilitate faster and more accurate generation of antibody-antigen complex structures. Structure-based design with antibody-specific diffusion models [37–39] may potentially deliver rational *de novo* biologics design against novel targets, although these models have seen limited experimental success to date, see e.g. Appendix E.

The findings of our study have practical implications for the development of therapeutic antibodies and pave the way for the computational design of developable therapeutic antibodies. By improving the efficiency and accuracy of antibody design, our pipeline has the potential to accelerate the development of new therapeutics and improve their success rates in clinical applications. We provide the code and data generated in this study under MIT license at https://github.com/Exscientia/ab-characterisation and https://doi.org/10.5281/zenodo.13862717.

## 4 Materials and Methods

### 4.1 Experimental methods

#### 4.1.1 Antibody protein production and purification

Protein production and gene synthesis were carried out by Genscript. After synthesis, selected VH/VL sequences were cloned onto an IgG1 backbone before transfection into a proprietary Genscript TurboCHOHT expression system. The purified, eluted protein fractions were then pooled and the buffer exchanged into PBS pH7.4. The final protein concentration was determined by the A280 method.

#### 4.1.2 Antigen protein production

GenScript provided pcDNA3.4 plasmids encoding the original Spike RBD variant as well as the XBB.1 and XBB.1.5 mutant strains with additional C-terminal hexahistidine (6xHis) tags for affinity purification. Plasmids were transiently transfected into Expi293F cells (ThermoFisher). The cellular density was adjusted to 3 *×* 10^6^ cell/mL, with a final volume of 400 mL of Expi293 expression media in 2-L non-baffled Erlenmeyer flasks (Corning, cat. No. 431255). Next, 400 mg of plasmid DNA was diluted with 24 mL of Opti-MEM medium (ThermoFisher, cat. No. 31985062). In a separate tube, 1.3 mL of Expifectamine 293 reagent (GIBCO, cat. No. 13385544) was then diluted in 23.7 mL of the same medium. Sample DNA and Expifectamine were then mixed and incubated at room temperature for 20 min. Cultured cells with this mixture added were then incubated for 3 days at 37°C and 8% CO2. The supernatant from the culture was collected by centrifugation at 4000xg for 30 min. Affinity purification was then performed using AmMag Ni Magnetic beads (GenScript Biotech, cat. No. U3108HC180). Beads were washed 4 times with 50 mM HEPES at pH 7.5, 500-mM NaCl, and 5% glycerol buffer to remove non-specifically bound protein. Protein was then eluted with 50-mM HEPES at pH 7.5, 500 mM NaCl, 5% glycerol, and 500 mM imidazole. Eluted Spike RBD was then buffer exchanged into 50-mM HEPES and 150-mM NaCl buffer with a PD-10 Sephadex G-25 desalting column (Cytiva, cat. No. 17085101). The purity and concentration of the obtained protein were determined by SDS-PAGE and the A280 method, respectively.

#### 4.1.3 Differential scanning fluorimetry experiments

Monoclonal antibody thermal stability was assessed with a Prometheus Panta instrument (NanoTemper Technologies) using a differential scanning fluorimetry (DSF) analysis with temperature increased from 25 to 95°C with a rate of increase of 1°C/min. High sensitivity capillaries supplied by NanoTemper Technologies (Cat. No. PR-C006) were used in all experiments. Output data were processed using a Prometheus Panta Control Software (NanoTemper Technologies). We report the lower melting temperature, Tm1, for all starting point and designed antibodies.

#### 4.1.4 Surface plasmon resonance experiments

Surface plasmon resonance (SPR) analysis of purified mAbs was performed with a Bruker Sierra Sensors MASS-2 instrument (Bruker Corp), following the metholody described in [40]. In these experiments, goat anti-human Fc antibody (Jackson ImmunoResearch, Product #109-005-088) was covalently coupled on an HCA sensorchip (Bruker) by first activating the spots with a mixture of 400 mM EDC and 100 mM NHS for 10 minutes at 10 µL/min. This step was followed by coupling of the anti-human Fc antibody at 25 µg/ml in sodium acetate buffer (10 mM, pH 5.0) for 6 minutes at 15 µL/min. The remaining uncoupled sites were blocked with 1 M ethanolamine at pH 8.0 injected for 6 minutes at 10 µL/min. In general, 13,000 RU of antibody was captured on each spot. After surface preparation, human mAb capture was performed at 20 µg/ml for 30 seconds at 10 µL/min to give on average 250 RU. Binding to three variants of SARS-CoV-2 RBD, the original strain, XBB.1 and XBB.1.5 mutants, was measured in independent experiments. For initial SPR experiments a single-shot injection of the three antigens was performed at 100 nM for 90 sec with a flow-rate of 50 µl/min and the dissociation phase was monitored for 600 sec. Confirmation of binders via titration experiments was performed with the same injection periods but with a 10-point three-fold serial dilution from 1 µM for each antigen. To regenerate the surface, 200 mM phosphoric acid was injected for 30 sec three times. The mAbs and antigens were diluted in the running buffer, which consisted of HBS-EP+ (10 mM HEPES pH 7.4, 150 mM NaCl, 3 mM EDTA, 0.05% v/v Tween-20). Data was double referenced by subtracting reference data obtained simultaneously with the active injection at a blank spot and subtracting data from a preceding blank buffer injection obtained at the active spot. Data processing was carried out with Sierra Analyser (Bruker, ver3.4.1) and fitting was carried out using a 1:1 model with Scrubber (BioLogic Software, ver2.0c).

#### 4.1.5 Surface plasmon resonance results analysis

The equilibrium dissociation constant *K_D_* is calculated from the ratio of the dissociation rate constant *k_d_* and the association rate constant *k_a_*, which are fitted from the corresponding slopes in the sensorgram obtained through the procedure described in Section 4.1.4. In practice, we report the negative logarithm of *K_D_*,

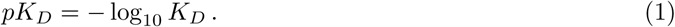

To enable easy comparison of binding results, we refer to binding detected in SPR experiments as weak (4.5 *< pK_D_ <* 6), medium (6 *≤ pK_D_ <* 8) or strong (8 *≤ pK_D_*). Further, for single concentration antigen SPR measurements, we introduce a quality assurance criterion, discounting any data points for which a binding signal was observed but which had a response of less than 30% of the theoretical maximum response as non-binding.

#### 4.1.6 Size-exclusion chromatography high-performance liquid chromatography experiments

Measurements were performed with an Agilent 1260 Infinity II system using AdvanceBioSEC 200A 1.9 µm 4.x6×150mm columns (Cat. No. PL1580-3201) and a PBS pH 7.4 mobile phase at a flow rate of 0.4ml/min for 7.2 minutes per injection. A diode array monitoring absorbance at 205 nm was used as the detector and peaks were integrated in the window of 1-4.1 minutes retention time. For each mAb, 100 ng of total protein were injected in a maximum injection volume of 10 *µ*l, with 10 *µ*l being injected for proteins that did not reach a purified concentration higher than 10ng/*µ*l. The fraction of the total integrated peak area contained within the highest peak was reported as “ratio in main peak”. Using a set of benchmark antibodies as reference, we considered antibody candidates as developable, if the highest peak was observed between two to three minutes retention time and has a ratio in the main peak above 0.5. This set of parameters was chosen to ensure that the highest peak was the monomeric peak and that the majority of the antibody was monomeric in solution. Note that here monomeric refers to the intact antibody protein in its native state. We use this metric to assess the aggregation state and hydrophobicity of antibodies [41].

### 4.2 Computational methods

#### 4.2.1 Antibody characterisation pipeline

In order to enable assessment, ranking and filtering of computationally suggested candidate antibodies prior to experimental validation, we implemented a computational pipeline to predict a range of metrics, both with and without the presence of the corresponding antigen, as shown in Figure 1. This pipeline is in the following referred to as the *characterisation pipeline*. The pipeline takes as input a set of antibody sequences to be characterised as well as a reference complex structure of a starting point antibody against the target of interest. Antibody structures and antibody-antigen complexes are predicted and potential liabilities (sequence- and structure-based) filtered out, prior to candidate ranking. To this end, antibody sequences are first annotated, characterised and filtered. For the remaining antibody sequences, structural models are predicted and assessed for developability as well as stability. Using target epitope and antibody structures, complex structures are predicted and the resulting complexes are ranked and filtered based on predicted interactions to determine candidate antibodies to take forward to experimental assessment.

##### Sequence-based characterisation

Antibody sequences are first numbered using ANARCI [42]. Candidate sequences are then scanned for any motifs that are known to negatively affect developability [43], and any that contain these liabilities are discarded at this stage in order to reduce downstream computational cost.

##### Structure-based antibody characterisation

Antibody structures are predicted using ABodyBuilder2 [44, 45]. Predicted structures are then characterised using with the Therapeutic Antibody Profiler (TAP) [46] and structures with red TAP flags are discarded from further analysis. In addition to this, interface stability of the heavy and light chains is characterised using Rosetta as described in section 4.2.6 and structures with predicted interface energy above 5 kcal/mol, used as a proxy for antibody instability, are discarded from downstream analysis.

##### Structure-based complex characterisation

The remaining candidate antibodies are characterised for their predicted interactions with the target antigen. ChimeraX [47] is used to generate a complex structure between each candidate antibody and the target antigen, using the antibody in the reference complex to guide the complex generation by calculating a low resolution map (6 Å) of the reference antibody and fitting the candidate antibody into that map. The resulting complex is relaxed using OpenMM [48].

Following complex generation, the predicted complex is characterised using Rosetta. Complexes with shape complementarity score below 0.45 and complexes with positive interaction energy at mutated residues (see section 4.2.6) are discarded at this point. The remaining structures are ranked as described below.

##### Ranking of antibody candidates

In order to balance overall structure stability and interface energetics, we employ a bi-variate Gaussian fit for the ranking of candidate antibodies after structure characterisation. To this end, we fit the bi-variate Gaussian to the overall Rosetta score and the Rosetta predicted free energy change. We omit from the fit any sequences which have either total score or free energy change further than 1.5 interquartile ranges from the upper or lower quartile boundaries. For designs where both the overall score and the free energy change are lower than the starting point antibody, we then compute the Mahalanobis distance, while all others are discarded. Designs are then ranked by decreasing distance.

##### Confirmation of high-ranking candidates

Rosetta calculations are stochastic, with multiple runs with the same input structure leading to an ensemble of results. Prior to experimental validation, we therefore repeat the Rosetta score generation for the 50 highest-ranking candidate antibodies with ten replicates. We then consider the two replicates with the lowest total score, as well as the two replicates with the lowest free energy change. Final Rosetta scores are calculated by averaging over this set of two to four replicates, and we perform an identical ranking procedure to select the designs with highest Mahalanobis distance in the left lower quadrant for experimental verification. Further details are given in Appendix D.

#### 4.2.2 Candidate library generation from OAS

We scanned both paired and unpaired OAS [22, 23] for potential candidates for assessment with the antibody characterisation pipeline outlined above. For paired OAS, where both light and heavy chain of each deposited antibody are known, we determined the V and J gene of the starting point antibody and considered all non-redundant, paired antibodies in OAS as candidates which had matching V and J gene in the heavy chain and a CDR H3 length within one residue of the starting antibody length. For unpaired OAS, we searched for heavy chains in OAS with the same V and J gene as the starting point antibody heavy chain, same length CDR H3 and at least 50% sequence identity to the starting point CDR H3. The heavy chains identified through this scan were paired with the starting point antibody light chain. Candidates are then assessed according to the characterisation pipeline described in Section 4.2.1. We further expanded the libraries through two rounds of elaboration using the inverse folding model described in section 4.2.3. In each round, we generated 25 variants for each of the 100 top ranked designs from each library, using the inverse folding model and assessed according to our characterisation pipeline.

#### 4.2.3 Candidate generation with an inverse folding model

To generate candidates with similar structural features but with novel sequences, we used the inverse folding model AbMPNN [29]. AbMPNN is a fine-tuned version of the ProteinMPNN [49] inverse folding model that specializes in antibody sequence prediction. The model takes an antibody-antigen complex structure as input and produces a sequence for the antibody that is predicted to fold into the complex structure. It can take partial sequence information for conditioning on both antibody and antigen sequence.

To select which residues to design, we identify all paratope residues in the CDR loops for the starting antibody structures. Here we define paratope residues as those with a heavy atom within 4 Å of an antigen heavy atom. We then selected the four CDR loops with the most paratope residues and redesigned the paratope residues in these loops. This was intended to reduce the edit distance to the starting antibody in comparison to modifying the entire paratope. We then generated 5,000 sequences per starting antibody at a temperature of T=0.2 for 25,000 sequences in total. Candidates are then assessed according to the characterisation pipeline described in Section 4.2.1.

#### 4.2.4 Candidate generation with a language model

We consider suggested mutations from the ESM-1b language model and the ESM-1v ensemble of five protein language models [24], following the language-model-guided affinity maturation procedure described in [25]. Amino acid substitutions along the heavy and light chain sequences are recommended by a consensus of masked language models, and we consider as potential mutations the set of substitutions with higher likelihood than the wildtype by at least one model.

We provide a list of all these potential substitutions along with the wildtype residue as input to Rosetta, allowing it to pick either a suggested language model mutation or the wildtype amino acid. Rosetta then samples rotamers to repack the side-chains of the modified antibody in complex with the antigen. We then use the same selection criteria as described in Section 4.2.1 to select the best scoring sequences from a sample of 1,000 sequences generated from combinations of all possible mutations, though only a single round of Rosetta calculations are performed in this pipeline due to the comparatively lower number of designs.

This leads to relatively high edit distances to the wildtype sequence, as Rosetta preferentially picks out some of the mutations consistently, and leads to diversity across samples due to the stochastic nature of the assessment.

#### 4.2.5 Generation of initial structures for design

Though a chimeric structure of XBB.1.5 RBD fused to the SARS-CoV-1 core structure has recently become available [50], no structure was available at the beginning of our study. To enable structure-based assessment of designs against XBB.1.5 RBD, we therefore predicted the structure of XBB.1.5 RBD using ColabFold [51]. The resulting model structure is predicted with high confidence except in the region proximal to the RBD glycan. This ColabFold structure was then aligned to the RBD models present in the template PDBs noted in Appendix B to generate antibody-XBB.1.5 RBD complexes. The resulting complexes were then energy relaxed with Rosetta as described in section 4.2.6.

#### 4.2.6 Physics-based minimization, design, and scoring with Rosetta

Rosetta3 [52] was used for relaxation and energetic scoring of antibody Fvs and antibody-antigen complexes. In the efficient evolution design pipelines, Rosetta was also used to introduce mutations into the CDRs of antibody-antigen complexes.

In all Rosetta runs the *beta nov16* score function was used to assess energies and the *InterfaceAnalyzeMover* used to compute *dG separated*, the Rosetta prediction of binding free energy change, at the interface of interest, given in kcal/mol. When the input to Rosetta was an antibody Fv, the interface between the heavy and light chains was analysed with *InterfaceAnalyzeMover*. In cases when an antibody-antigen complex was input to Rosetta, interface analysis was applied to the antibody-antigen interface. In all cases, side chains were repacked before computing interface statistics. The *RunSimpleMetrics* [53] function was used to compute the structural aggregation propensity (SAP) [54], RMSD from the initial structure, and dihedral distance from the initial structure. We briefly summarize the key Rosetta protocols used in our study below.

In relaxation simulations, all backbone atom positions were restrained with *weights* = 1 and the structure then minimised to an energy tolerance of 0.001 kcal/mol using *MinMover*. This minimised structure was then subjected to one iteration of the *FastRelax* repacking and minimization protocol, and the resulting structure then assessed with *RunSimpleMetrics* and *InterfaceAnalyzeMover*.

For design, Rosetta format resfiles were prepared specifying which amino acid positions were mutable and to which new amino acids they could change. First, restraints were applied to all *C_α_* atoms in the input structure to maintain the overall backbone configuration. Mutations are introduced and the structure relaxed via three alternating rounds of rotamer repacking and minimization with soft constraints (*weights* = 0.4) using *PackRotamersMover* and *MinMover*, respectively, followed by two rounds of minimization with *RotamersTrialsMinMover*. Each design cycle was concluded by a final minimisation with *MinMover*. This process was repeated three times and the final structure analysed with *InterfaceAnalyzeMover* and *RunSimpleMetrics* as noted above.

Due to the stochastic nature of the Rosetta relaxation and design protocols, repeat runs using the same starting structure give different results. We therefore carried out ten replicate runs of Rosetta and considered the set of up to four replicate structures with the two lowest total score values and the two lowest free energy change to compute final scoring metrics.

We also used the *interface energy* module within Rosetta to predict the degree to which antibodies engage SARS-CoV-2 XBB.1.5 RBD. First, we used PyMol to list the paratope residues in the antibody as those within 4 ^Å^ of the antigen and the epitope residues in the antigen as those within 4 ^Å^ of the antibody. We then computed all pairwise contact energies between these two sets of residues to generate a contact energy matrix. Finally, we computed the sum of all pairwise contact energies involving any of the 23 XBB.1.5 RBD mutations, generating a single energy value representing the strength of interactions of the antibody with mutated residues in XBB.1.5 RBD. We exclude any candidates for which the difference between the sum of mutated energies is positive.

### 4.3 Cryogenic electron microscopy methods

#### 4.3.1 Sample preparation

Purified Fabs were incubated with 6P stabilised SARS-CoV-2 spike protein [55] at a molar ratio of 1.2:1 for 5 minutes on ice. Prior to application of this solution to the grid, fluorinated octyl maltoside (FOM) was added to a final concentration of 0.01% (w/v) to aid in overcoming preferred orientation. In all cases 3 µl of spike:Fab complex solution was applied to Quantifoil R1.2/1.3 grids (200 mesh) pre-treated by glow discharge in air at 20 mAmp for 30 seconds, resulting in a negatively charged grid surface. Samples were blotted with 0 blot force for 4 seconds, and plunge frozen in liquid ethane using a Vitrobot Mark IV (Thermo Fisher Scientific). The sample chamber was held at 4 °C and 95% relative humidity throughout.

#### 4.3.2 Data collection

Cryo-EM data were collected at the Eindhoven NanoPort facility on a 200 kV Glacios 2 cryo-TEM (Thermo Fisher Scientific) equipped with a Falcon 4i direct electron detector and Selectris energy filter. Further details on optical settings are listed in Appendix C.

#### 4.3.3 Image processing

Cryo-EM image processing was carried out in cryoSPARC 4.0 [56]. Raw EER movies were preprocessed using patch motion correction and patch CTF estimation without upsampling. Particles were picked with cryoSPARC blob picker and extracted into 520 pixel boxes, binned to 130 for early processing steps. Iterative 2D classification and manual curation were used to clean the initial particle stack. *Ab initio* model generation into 4 classes was used to distinguish spike:fab complexes from apo spike. Particles containing spike:fab complex were re-extracted into 2x bin (260 px) boxes for further refinements. Nonuniform refinement was used to generate global 3D reconstructions with C1 symmetry. Local refinement masks comprising RBD and Fab were generated with UCSF Chimera [47] and used for local refinement of spike-fab interfaces in cryoSPARC. Final maps were locally sharpened with deepEMhancer [57]. FSCs were calculated using the gold-standard method at a cut-off of 0.143.

## Acknowledgements

We thank John Overington, Andrew Hopkins, Jody Barbeau, Eileen Brandenburger, Anthony Bradley, Sean Robinson, Tjelvar Olsson, Luis Yanes and Iva Hopkins Navratilova for useful discussions and comments. We also thank Berend Jan Bosch and Chunyan Wang from Utrecht University for providing the SARS-CoV-2 spike protein used in this study.

## Appendix A Class 3 antibody SPR screen

We performed a broad single concentration antigen screen with the SPR method described in Section 4.1.4 to estimate binding affinity of 192 class 3 antibodies against the Wuhan strain, the XBB.1 strain and the XBB.1.5 strain. A summary of these measurements is given in Table A1, with further details available in the full dataset at https://doi.org/10.5281/zenodo.13862717.

**Table A1.**
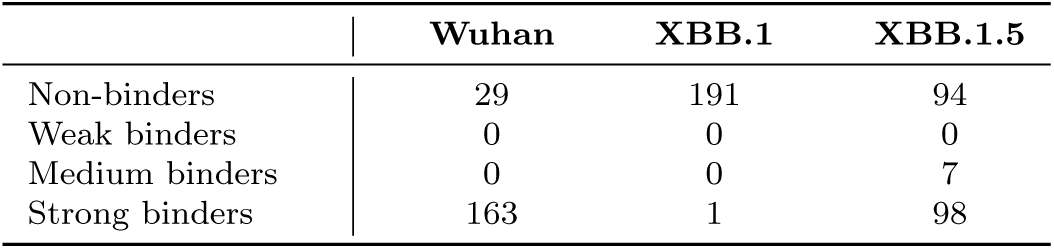
Summary of experimental characterisation of 192 class 3 antibodies from a single concentration antigen screen against three strains of the SARS-CoV-2 RBD. Binding strength is determined from the *pK_D_* value as detailed in Section 4.1.5.

Using the cryogenic electron microscopy methodology described in Section 4.3, we assess the binding pose and antibody structure of two selected sequences, OC220225-SN0108 and OC220302-SN0554. These are from the set of 182 for which no previously known structure exists and exhibit strong binding to the Wuhan strain. The structural characterisation for both antibodies is shown in Figure A1.

**Fig. A1.**
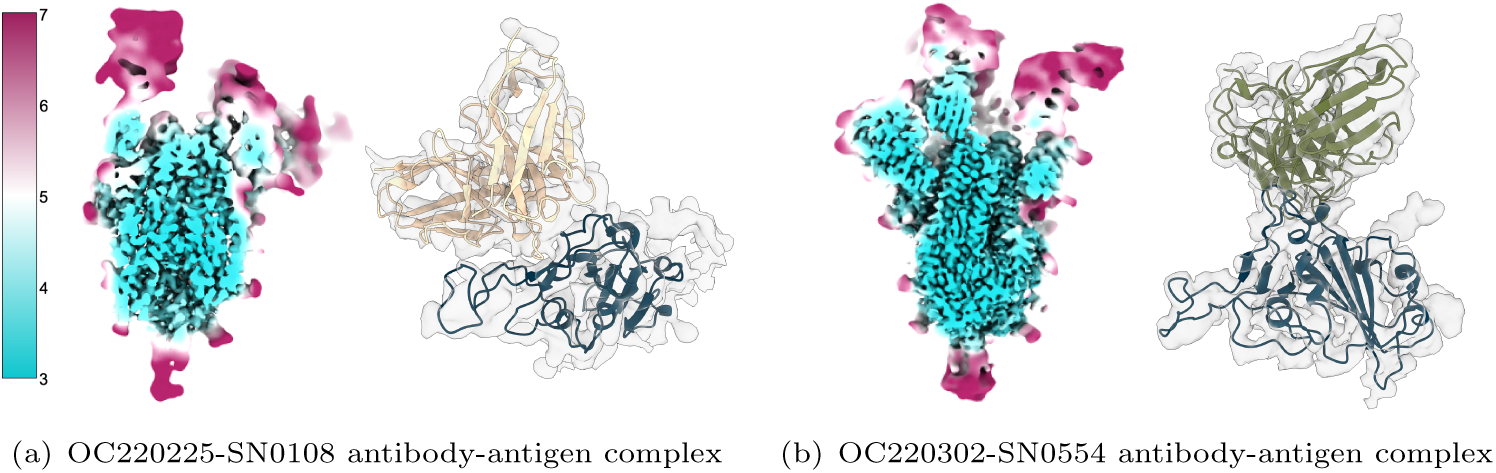
Cryo-electron microscopy epitope mapping of two selected antibodies in complex with the SARS-CoV-2 spike protein. In both panels, the left-hand side shows central slices of 3D electron microscopy volumes coloured by local resolution in angstrom, and the right-hand side displays locally refined interfaces of the fragment antigen-binding region and receptor-binding domain (in blue).

## Appendix B Starting point antibodies

The starting point antibodies used in this article are shown in Table B2. Of these six starting points, all have potent binding to the Wuhan strain of the virus, but only S309 is an effective binder to all three antigens tested, namely wild-type, XBB.1 and XBB.1.5. S2K146, BD55-5840 and DXP-604 are weak binders to XBB.1.5, but do not bind to XBB.1. REGN10987 and LY-CoV1404 only bind to the Wuhan strain and have no efficacy against XBB.1 and XBB.1.5.

**Table B2.** Starting point antibodies used in this study, along with the template PDB structure used, and the corresponding RBD strain. Each starting point was used with the Efficient Evolution approach, while only starting point antibodies with CDR H3 of length up to 15 residues were used in the inverse folding and OAS scanning method.

| Antibody Name | Template PDB used | RBD Strain in template PDB | Used in method |  |  |
| --- | --- | --- | --- | --- | --- |
|  |  |  | *Inverse folding* | *OAS scanning* | *Efficient Evol.* |
| BD55-5840 | 7WRZ [58] | Omicron | ✓ | ✓ | ✓ |
| DXP-604 | 7CH4 [59] | Wuhan | ✓ | ✓ | ✓ |
| LY-CoV1404 | 7MMO [60] | Wuhan | ✓ | ✓ | ✓ |
| REGN10987 | 6XDG [61] | Wuhan | ✓ | ✓ | ✓ |
| S2K146 | 7TAS [62] | Wuhan | ✓ | ✓ | ✓ |
| S309 | 7X1M [63] | Omicron | — | — | ✓ |

## Appendix C Cryo-EM settings

Details on optical settings for the cryo-EM data collection are given in Table C3.

**Table C3.** Data collection parameters.

| Samples | REGN10987 antibody- antigen complex<br>S309 antibody-antigen complex | Design derived from the DXP-604 starting point<br>Design derived from the LY-CoV1404 starting point<br>OC220225-SN0108 antibody-antigen complex<br>OC220302-SN0554 antibody-antigen complex |
| --- | --- | --- |
| Energy source | XFEG | E-CFEG |
| Voltage (kV) | 200 | 200 |
| Magnification | 165,000 | 165,000 |
| Apertures | C2: 50 $\mu m$, no objective aperture | C2: 30 $\mu m$, no objective aperture |
| Spot size | 2 | 2 |
| Beam diameter ($\mu m$) | 1.1 | 1.1 |
| Pixel size ($\text{\AA}$) | 0.694 | 0.694 |
| Dose rate (e-/pix/sec) | 10.8 | 12.1 |
| Electron exposure (e-/$\text{\AA}^2$) | 50 | 50 |
| EER frames | 702 | 657 |
| Exposure time (sec) | 2.29 | 2.15 |
| Selectris energy filter slit (eV) | 10 | 10 |
| Images per hole | 2 (Fringe-Free Imaging) | 2 (Fringe-Free Imaging) |
| Hole centering | Aberration-free image shift | Aberration-free image shift |
| Delay after stage shift (sec) | 10 | 5 |
| Delay after image shift (sec) | 0.5 | 0.5 |
| Autofocus | Once per cluster | Once per cluster |
| Defocus range ($\mu m$) | -0.7 to -1.4 | -0.7 to -1.4 |

## Appendix D Ranking of designs

In Figure D2, we show the distribution of total score and free energy change for all candidates across each design strategy. For the language model pipeline, which had a smaller number of total designs, ten replicates were used in a single round to select final top-ranking candidates. An inset is shown for each plot, displaying the left lower quadrant of the distribution, with contour plots of the bi-variate Gaussian fit used for final ranking according to Mahalanobis distance. The fit excludes outliers, defined as further than 1.5 interquartile range above the third quartile or below the first quartile.

**Fig. D2.**
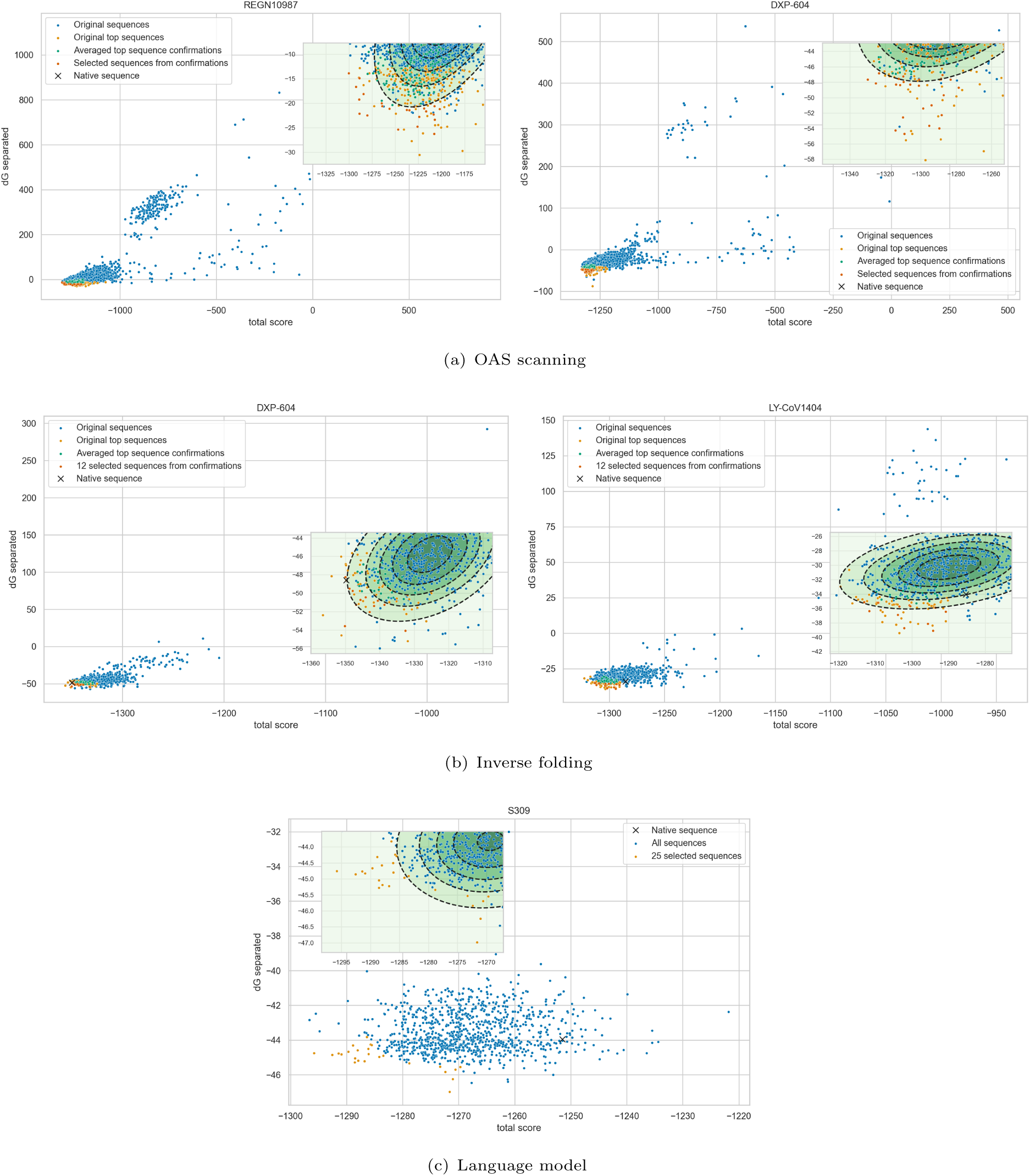
Multi-variate gaussian fit of the designed candidates, with selected designs shown in orange. (a) Two rounds of Rosetta selection for the OAS scanning pipeline, with one and ten replicates respectively. (b) Two rounds of Rosetta selection for the inverse folding pipeline, with one and ten replicates respectively. (c) Selection of language model designs from the efficient evolution pipeline.

## Appendix E CDR design with RFdiffusion

We attempted to use RFdiffusion [64], a state-of-the-art protein backbone diffusion model, to re-design CDR loops of starting point antibodies. We considered two starting point antibodies, BD55-5840 and LY-Cov1404, and used RFdiffusion to design part of the CDR H3 and CDR L3 loops respectively, using the remainder of the antibody and the XBB.1.5 antigen as motif.

We identify the six contiguous residues closest to the antigen as those to re-design in each selected CDR loop, and sample backbone conformations with RFdiffusion of six to nine residues to replace the original CDR loop. We use four different configurations for both starting point: (1) the default RFdiffusion model, (2) the Complex beta ckpt model weights, intended to generate a greater diversity of backbone topologies, (3) partially noising and de-noise the original backbone with 20 steps with the default model weights, and (4) using the default model in conjunction with an auxiliary potential, which consists of an SE(3) transformer trained on classifying experimental antibody structures from modified ones where all CDR loops have been re-designed with RFdiffusion. For each configuration and starting point, we generate 200 samples, for a total of 800 structures. For each structure, we then generate 5 sequences by predicting all modified residues with AbMPNN.

Across each pool of 1000 sequences and backbone structures for every configuration and starting point, we select 10 using the characterisation pipeline described in Section 4.2.1 for experimental validation. These 80 designs, 40 for each starting point, are selected to bind against the XBB.1.5 variant. As shown in Table E4, we find that none of these designs bind to XBB.1 or XBB.1.5, however 56 of them show efficacy against the Wuhan strain, to which the starting points also had measurable binding, resulting in a hit rate of 70%. There is no statistically significant difference in measured binding affinity or hit rate across the different configurations. Of the 56 binders, 25 have modified loop lengths where additional residues where inserted during the CDR backbone redesign.

**Table E4.**
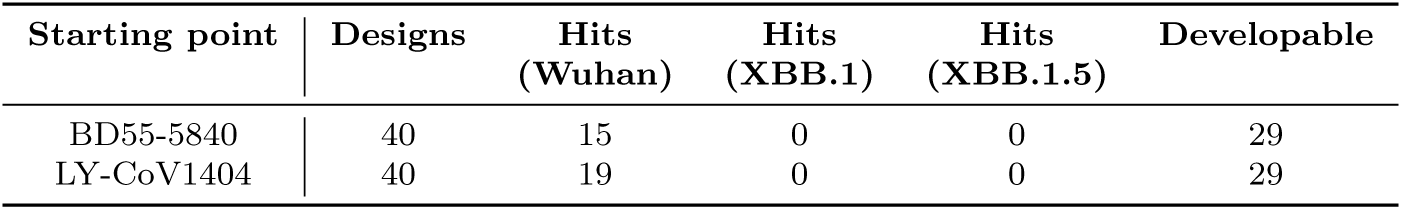
Summary of experimental characterisation of candidate antibodies generated with RFdiffusion.

| Starting point | Designs | Hits<br>(Wuhan) | Hits<br>(XBB.1) | Hits<br>(XBB.1.5) | Developable |
| --- | --- | --- | --- | --- | --- |
| BD55-5840 | 40 | 15 | 0 | 0 | 29 |
| LY-CoV1404 | 40 | 19 | 0 | 0 | 29 |

## References

[1] Crescioli, S., Kaplon, H., Chenoweth, A., Wang, L., Visweswaraiah, J., Reichert, J.M.: Antibodies to watch in 2024. mAbs 16(1), 2297450 (2024) 10.1080/19420862.2023.2297450. PMID: 38178784

[2] Walsh, G., Walsh, E.: Biopharmaceutical benchmarks 2022. Nature Biotechnology 40(12), 1722–1760 (2022)

[3] Mullard, A.: Fda approves 100th monoclonal antibody product. Nature reviews. Drug discovery 20(7), 491–495 (2021)

[4] https://www.pharmaceutical-technology.com/analyst-comment/biologic-sales-small-molecule-sales/

[5] Kim, J., McFee, M., Fang, Q., Abdin, O., Kim, P.M.: Computational and artificial intelligence-based methods for antibody development. Trends in Pharmacological Sciences 44(3), 175–189 (2023)

[6] Silva, B.M., Myung, Y., Ascher, D.B., Pires, D.E.V.: epitope3D: a machine learning method for conformational B-cell epitope prediction. Briefings in Bioinformatics 23(1), 423 (2021) 10.1093/bib/bbab423 https://academic.oup.com/bib/article-pdf/23/1/bbab423/42230615/bbab423.pdf

[7] Chinery, L., Wahome, N., Moal, I., Deane, C.M.: Paragraph—antibody paratope prediction using graph neural networks with minimal feature vectors. Bioinformatics 39(1), 732 (2022) 10.1093/bioinformatics/btac732 https://academic.oup.com/bioinformatics/article-pdf/39/1/btac732/48448850/btac732.pdf

[8] Kurumida, Y., Saito, Y., Kameda, T.: Predicting antibody affinity changes upon mutations by combining multiple predictors. Scientific reports 10(1), 19533 (2020)

[9] Adolf-Bryfogle, J., Kalyuzhniy, O., Kubitz, M., Weitzner, B.D., Hu, X., Adachi, Y., Schief, W.R., Dunbrack Jr, R.L.: Rosettaantibodydesign (rabd): A general framework for computational antibody design. PLoS computational biology 14(4), 1006112 (2018)

[10] Zhang, W., Wang, H., Feng, N., Li, Y., Gu, J., Wang, Z.: Developability assessment at early-stage discovery to enable development of antibody-derived therapeutics. Antibody Therapeutics 6(1), 13–29 (2023)

[11] Bachas, S., Rakocevic, G., Spencer, D., Sastry, A.V., Haile, R., Sutton, J.M., Kasun, G., Stachyra, A., Gutierrez, J.M., Yassine, E., et al.: Antibody optimization enabled by artificial intelligence predictions of binding affinity and naturalness. BioRxiv, 2022–08 (2022)

[12] Gruver, N., Stanton, S., Frey, N., Rudner, T.G., Hotzel, I., Lafrance-Vanasse, J., Rajpal, A., Cho, K., Wilson, A.G.: Protein design with guided discrete diffusion. Advances in Neural Information Processing Systems 36 (2024)

[13] Shanker, V.R., Bruun, T.U., Hie, B.L., Kim, P.S.: Inverse folding of protein complexes with a structure-informed language model enables unsupervised antibody evolution. bioRxiv, 2023–12 (2023)

[14] Hummer, A.M., Abanades, B., Deane, C.M.: Advances in computational structure-based antibody design. Current Opinion in Structural Biology 74, 102379 (2022)

[15] Shanehsazzadeh, A., McPartlon, M., Kasun, G., Steiger, A.K., Sutton, J.M., Yassine, E., McCloskey, C., Haile, R., Shuai, R., Alverio, J., Rakocevic, G., Levine, S., Cejovic, J., Gutierrez, J.M., Morehead, A., Dubrovskyi, O., Chung, C., Luton, B.K., Diaz, N., Kohnert, C., Consbruck, R., Carter, H., LaCombe, C., Bist, I., Vilaychack, P., Anderson, Z., Xiu, L., Bringas, P., Alarcon, K., Knight, B., Radach, M., Bateman, K., Kopec-Belliveau, G., Chapman, D., Bennett, J., Ventura, A.B., Canales, G.M., Gowda, M., Jackson, K.A., Caguiat, R., Brown, A., Silva, D.G., Guo, Z., Abdulhaqq, S., Klug, L.R., Gander, M., Yapici, E., Meier, J., Bachas, S.: Unlocking de novo antibody design with generative artificial intelligence. bioRxiv (2024) 10.1101/2023.01.08.523187 https://www.biorxiv.org/content/early/2024/01/07/2023.01.08.523187.full.pdf

[16] Cheng, J., Liang, T., Xie, X.-Q., Feng, Z., Meng, L.: A new era of antibody discovery: an in-depth review of ai-driven approaches. Drug Discovery Today 29(6), 103984 (2024) 10.1016/j.drudis.2024.103984

[17] Kim, J., McFee, M., Fang, Q., Abdin, O., Kim, P.: Computational and artificial intelligence-based methods for antibody development. Trends in Pharmacological Sciences 44 (2023) 10.1016/j.tips.2022.12.005

[18] Svilenov, H.L., Arosio, P., Menzen, T., Tessier, P., Sormanni, P.: Approaches to expand the conventional toolbox for discovery and selection of antibodies with drug-like physicochemical properties 15(1), 2164459 (2023). Taylor & Francis

[19] Nimrod, G., Fischman, S., Austin, M., Herman, A., Keyes, F., Leiderman, O., Hargreaves, D., Strajbl, M., Breed, J., Klompus, S., Minton, K., Spooner, J., Buchanan, A., Vaughan, T.J., Ofran, Y.: Computational design of epitope-specific functional antibodies. Cell reports 25(8), 2121–21315 (2018) 10.1016/j.celrep.2018.10.081

[20] Jette, C.A., Cohen, A.A., Gnanapragasam, P.N.P., Muecksch, F., Lee, Y.E., Huey-Tubman, K.E., Schmidt, F., Hatziioannou, T., Bieniasz, P.D., Nussenzweig, M.C., West, A.P., Keeffe, J.R., Bjorkman, P.J., Barnes, C.O.: Broad cross-reactivity across sarbecoviruses exhibited by a subset of covid-19 donor-derived neutralizing antibodies. Cell Reports 36(13), 109760 (2021) 10.1016/j.celrep.2021.109760

[21] Cao, Y., Wang, J., Jian, F., Xiao, T., Song, W., Yisimayi, A., Huang, W., Li, Q., Wang, P., An, R., Wang, Y., Niu, X., Yang, S., Liang, H., Sun, H., Li, T., Yu, Y., Cui, Q., Liu, S., Yang, X., Du, S., Zhang, Z., Hao, X., Shao, F., Jin, R., Wang, X., Xiao, J., Wang, Y., Xie, X.S.: Omicron escapes the majority of existing sars-cov-2 neutralizing antibodies. Nature 602(7898), 657–663 (2022) 10.1038/s41586-021-04385-3

[22] Olsen, T.H., Boyles, F., Deane, C.M.: Observed antibody space: A diverse database of cleaned, annotated, and translated unpaired and paired antibody sequences. Protein Science 31(1), 141–146 (2022)

[23] Olsen, T.H., Boyles, F., Deane, C.M.: Observed antibody space: A diverse database of cleaned, annotated, and translated unpaired and paired antibody sequences. Protein Science 31(1), 141–146 (2022) 10.1002/pro.4205 https://onlinelibrary.wiley.com/doi/pdf/10.1002/pro.4205

[24] Rives, A., Meier, J., Sercu, T., Goyal, S., Lin, Z., Liu, J., Guo, D., Ott, M., Zitnick, C.L., Ma, J., Fergus, R.: Biological structure and function emerge from scaling unsupervised learning to 250 million protein sequences. Proceedings of the National Academy of Sciences 118(15), 2016239118 (2021) 10.1073/pnas.2016239118 https://www.pnas.org/doi/pdf/10.1073/pnas.2016239118

[25] Hie, B.L., Shanker, V.R., Xu, D., Bruun, T.U., Weidenbacher, P.A., Tang, S., Wu, W., Pak, J.E., Kim, P.S.: Efficient evolution of human antibodies from general protein language models. Nature Biotechnology 42(2), 275–283 (2024)

[26] Pham, N.B., Meng, W.S.: Protein aggregation and immunogenicity of biotherapeutics. International Journal of Pharmaceutics 585, 119523 (2020) 10.1016/j.ijpharm.2020.119523

[27] Greaney, A.J., Starr, T.N., Barnes, C.O., Weisblum, Y., Schmidt, F., Caskey, M., Gaebler, C., Cho, A., Agudelo, M., Finkin, S., Wang, Z., Poston, D., Muecksch, F., Hatziioannou, T., Bieniasz, P.D., Robbiani, D.F., Nussenzweig, M.C., Bjorkman, P.J., Bloom, J.D.: Mapping mutations to the sarscov-2 rbd that escape binding by different classes of antibodies. Nature Communications 12(1), 4196 (2021) 10.1038/s41467-021-24435-8

[28] Desautels, T.A., Arrildt, K.T., Zemla, A.T., Lau, E.Y., Zhu, F., Ricci, D., Cronin, S., Zost, S.J., Binshtein, E., Scheaffer, S.M., Dadonaite, B., Petersen, B.K., Engdahl, T.B., Chen, E., Handal, L.S., Hall, L., Goforth, J.W., Vashchenko, D., Nguyen, S., Weilhammer, D.R., Lo, J.K.-Y., Rubinfeld, B., Saada, E.A., Weisenberger, T., Lee, T.-H., Whitener, B., Case, J.B., Ladd, A., Silva, M.S., Haluska, R.M., Grzesiak, E.A., Earnhart, C.G., Hopkins, S., Bates, T.W., Thackray, L.B., Segelke, B.W., Lyon, E.Z.A., Anderson, P.S., Avila-Herrera, A., Bennett, W.F., Bourguet, F.A., Chen, J.C., Coleman, M.A., Collette, N.M., Davis, A., Vannest, B.D., Fong, E.J., Gilmore, S., Goncalves, A.R., Hall, S.B., Harmon, B., He, W., Hoang-Phou, S.A., Landajuela, M., Laurence, T.A., Lee, T.H., Da Silva, F.L., Liu, C., Mundhenk, T.N., Mohagheghi, M.V., McIlroy, P.R., Pham, L.T.M., Sanchez, J.C., Sinha, A., Solomon, E.A., Watkins, N., Yang, J., Ye, C., Zhang, B., Lillo, A.M., Sundaram, S., Bloom, J.D., Diamond, M.S., Crowe, J.E., Carnahan, R.H., Faissol, D.M., Consortium, T.-l.C.-.: Computationally restoring the potency of a clinical antibody against omicron. Nature 629(8013), 878–885 (2024) 10.1038/s41586-024-07385-1

[29] Dreyer, F.A., Cutting, D., Schneider, C., Kenlay, H., Deane, C.M.: Inverse folding for antibody sequence design using deep learning. The 2023 ICML Workshop on Computational Biology (2023) https://arxiv.org/abs/2310.19513

[30] Yang, Z., Milas, K.A., White, A.D.: Now what sequence? pre-trained ensembles for bayesian optimization of protein sequences. bioRxiv (2022) 10.1101/2022.08.05.502972 https://www.biorxiv.org/content/early/2022/08/06/2022.08.05.502972.full.pdf

[31] Stanton, S., Maddox, W., Gruver, N., Maffettone, P., Delaney, E., Greenside, P., Wilson, A.G.: Accelerating Bayesian Optimization for Biological Sequence Design with Denoising Autoencoders (2022)

[32] Khan, A., Cowen-Rivers, A.I., Grosnit, A., Deik, D.-G.-X., Robert, P.A., Greiff, V., Smorodina, E., Rawat, P., Dreczkowski, K., Akbar, R., Tutunov, R., Bou-Ammar, D., Wang, J., Storkey, A., Bou-Ammar, H.: AntBO: Towards Real-World Automated Antibody Design with Combinatorial Bayesian Optimisation (2022)

[33] Park, J.W., Stanton, S., Saremi, S., Watkins, A., Dwyer, H., Gligorijevic, V., Bonneau, R., Ra, S., Cho, K.: PropertyDAG: Multi-objective Bayesian optimization of partially ordered, mixed-variable properties for biological sequence design (2022)

[34] Park, J.W., Tagasovska, N., Maser, M., Ra, S., Cho, K.: BOtied: Multi-objective Bayesian optimization with tied multivariate ranks (2024)

[35] Abramson, J., Adler, J., Dunger, J., Evans, R., Green, T., Pritzel, A., Ronneberger, O., Willmore, L., Ballard, A.J., Bambrick, J., Bodenstein, S.W., Evans, D.A., Hung, C.-C., O’Neill, M., Reiman, D., Tunyasuvunakool, K., Wu, Z., Žemgulytė, A., Arvaniti, E., Beattie, C., Bertolli, O., Bridgland, A., Cherepanov, A., Congreve, M., Cowen-Rivers, A.I., Cowie, A., Figurnov, M., Fuchs, F.B., Gladman, H., Jain, R., Khan, Y.A., Low, C.M.R., Perlin, K., Potapenko, A., Savy, P., Singh, S., Stecula, A., Thillaisundaram, A., Tong, C., Yakneen, S., Zhong, E.D., Zielinski, M., Žídek, A., Bapst, V., Kohli, P., Jaderberg, M., Hassabis, D., Jumper, J.M.: Accurate structure prediction of biomolecular interactions with alphafold 3. Nature (2024) 10.1038/s41586-024-07487-w

[36] Ketata, M.A., Laue, C., Mammadov, R., Stärk, H., Wu, M., Corso, G., Marquet, C., Barzilay, R., Jaakkola, T.S.: Diffdock-pp: Rigid protein-protein docking with diffusion models. arXiv preprint arXiv:2304.03889 (2023)

[37] Martinkus, K., Ludwiczak, J., Cho, K., Liang, W.-C., Lafrance-Vanasse, J., Hotzel, I., Rajpal, A., Wu, Y., Bonneau, R., Gligorijevic, V., Loukas, A.: AbDiffuser: Full-Atom Generation of in vitro Functioning Antibodies (2024)

[38] Bennett, N.R., Watson, J.L., Ragotte, R.J., Borst, A.J., See, D.L., Weidle, C., Biswas, R., Shrock, E.L., Leung, P.J.Y., Huang, B., Goreshnik, I., Ault, R., Carr, K.D., Singer, B., Criswell, C., Vafeados, D., Garcia Sanchez, M., Kim, H.M., Vázquez Torres, S., Chan, S., Baker, D.: Atomically accurate de novo design of single-domain antibodies. bioRxiv (2024) 10.1101/2024.03.14.585103 https://www.biorxiv.org/content/early/2024/03/18/2024.03.14.585103.full.pdf

[39] Cutting, D., Dreyer, F.A., Errington, D., Schneider, C., Deane, C.M.: De novo antibody design with SE(3) diffusion (2024). https://arxiv.org/abs/2405.07622

[40] Kamat, V., Rafique, A., Huang, T., Olsen, O., Olson, W.: The impact of different human igg capture molecules on the kinetics analysis of antibody-antigen interaction. Analytical Biochemistry 593, 113580 (2020) 10.1016/j.ab.2020.113580

[41] Bailly, M., Mieczkowski, C., Juan, V., Metwally, E., Tomazela, D., Baker, J., Uchida, M., Kofman, E., Raoufi, F., Motlagh, S., Yu, Y., Park, J., Raghava, S., Welsh, J., Rauscher, M., Raghunathan, G., Hsieh, M., Chen, Y.-L., Nguyen, H.T., Nguyen, N., Cipriano, D., Fayadat-Dilman, L.: Predicting antibody developability profiles through early stage discovery screening. mAbs 12(1), 1743053 (2020) 10.1080/19420862.2020.1743053 10.1080/19420862.2020.1743053. PMID: 32249670

[42] Dunbar, J., Deane, C.M.: Anarci: antigen receptor numbering and receptor classification. Bioinformatics 32(2), 298–300 (2016)

[43] Leem, J., Dunbar, J., Georges, G., Shi, J., Deane, C.M.: ABodyBuilder: Automated antibody structure prediction with data–driven accuracy estimation. mAbs 8(7), 1259–1268 (2016) 10.1080/19420862.2016.1205773

[44] Abanades, B., Wong, W.K., Boyles, F., Georges, G., Bujotzek, A., Deane, C.M.: Immunebuilder: Deep-learning models for predicting the structures of immune proteins. Communications Biology 6(1), 575 (2023)

[45] Kenlay, H., Dreyer, F.A., Cutting, D., Nissley, D., Deane, C.M.: ABodyBuilder3: Improved and scalable antibody structure predictions (2024)

[46] Raybould, M.I., Deane, C.M.: The therapeutic antibody profiler for computational developability assessment. Therapeutic Antibodies: Methods and Protocols, 115–125 (2022)

[47] Goddard, T.D., Huang, C.C., Meng, E.C., Pettersen, E.F., Couch, G.S., Morris, J.H., Ferrin, T.E.: Ucsf chimerax: Meeting modern challenges in visualization and analysis. Protein science 27(1), 14–25 (2018)

[48] Eastman, P., Swails, J., Chodera, J.D., McGibbon, R.T., Zhao, Y., Beauchamp, K.A., Wang, L.- P., Simmonett, A.C., Harrigan, M.P., Stern, C.D., et al.: Openmm 7: Rapid development of high performance algorithms for molecular dynamics. PLoS computational biology 13(7), 1005659 (2017)

[49] Dauparas, J., Anishchenko, I., Bennett, N., Bai, H., Ragotte, R.J., Milles, L.F., Wicky, B.I.M., Courbet, A., Haas, R.J., Bethel, N., Leung, P.J.Y., Huddy, T.F., Pellock, S., Tischer, D., Chan, F., Koepnick, B., Nguyen, H., Kang, A., Sankaran, B., Bera, A.K., King, N.P., Baker, D.: Robust deep learning–based protein sequence design using proteinmpnn. Science 378(6615), 49–56 (2022) 10.1126/science.add2187 https://www.science.org/doi/pdf/10.1126/science.add2187

[50] Zhang, W., Shi, K., Geng, Q., Herbst, M., Wang, M., Huang, L., Bu, F., Liu, B., Aihara, H., Li, F.: Structural evolution of sars-cov-2 omicron in human receptor recognition. Journal of Virology 97(8), 00822–23 (2023) 10.1128/jvi.00822-23 https://journals.asm.org/doi/pdf/10.1128/jvi.00822-23

[51] Mirdita, M., Schütze, K., Moriwaki, Y., Heo, L., Ovchinnikov, S., Steinegger, M.: Colabfold: making protein folding accessible to all. Nature methods 19(6), 679–682 (2022)

[52] Leaver-Fay, A., Tyka, M., Lewis, S.M., Lange, O.F., Thompson, J., Jacak, R., Kaufman, K.W., Renfrew, P.D., Smith, C.A., Sheffler, W., Davis, I.W., Cooper, S., Treuille, A., Mandell, D.J., Richter, F., Ban, Y.-E.A., Fleishman, S.J., Corn, J.E., Kim, D.E., Lyskov, S., Berrondo, M., Mentzer, S., Popović, Z., Havranek, J.J., Karanicolas, J., Das, R., Meiler, J., Kortemme, T., Gray, J.J., Kuhlman, B., Baker, D., Bradley, P.: Chapter nineteen - rosetta3: An object-oriented software suite for the simulation and design of macromolecules. In: Johnson, M.L., Brand, L. (eds.) Computer Methods, Part C. Methods in Enzymology, vol. 487, pp. 545–574. Academic Press,(2011). 10.1016/B978-0-12-381270-4.00019-6 . https://www.sciencedirect.com/science/article/pii/B9780123812704000196

[53] Adolf-Bryfogle, J., Labonte, J.W., Kraft, J.C., Shapovalov, M., Raemisch, S., Lütteke, T., DiMaio, F., Bahl, C.D., Pallesen, J., King, N.P., Gray, J.J., Kulp, D.W., Schief, W.R.: Growing glycans in rosetta: Accurate de novo glycan modeling, density fitting, and rational sequon design. bioRxiv (2021) 10.1101/2021.09.27.462000 https://www.biorxiv.org/content/early/2021/10/08/2021.09.27.462000.full.pdf

[54] Lauer, T.M., Agrawal, N.J., Chennamsetty, N., Egodage, K., Helk, B., Trout, B.L.: Developability index: A rapid in silico tool for the screening of antibody aggregation propensity. Journal of Pharmaceutical Sciences 101(1), 102–115 (2012) 10.1002/jps.22758

[55] Hsieh, C.-L., Goldsmith, J.A., Schaub, J.M., DiVenere, A.M., Kuo, H.-C., Javanmardi, K., Le, K.C., Wrapp, D., Lee, A.G., Liu, Y., Chou, C.-W., Byrne, P.O., Hjorth, C.K., Johnson, N.V., Ludes-Meyers, J., Nguyen, A.W., Park, J., Wang, N., Amengor, D., Lavinder, J.J., Ippolito, G.C., Maynard, J.A., Finkelstein, I.J., McLellan, J.S.: Structure-based design of prefusion-stabilized sars-cov-2 spikes. Science 369(6510), 1501–1505 (2020) 10.1126/science.abd0826 https://www.science.org/doi/pdf/10.1126/science.abd0826

[56] Punjani, A., Rubinstein, J.L., Fleet, D.J., Brubaker, M.A.: cryoSPARC: algorithms for rapid unsupervised cryo-EM structure determination. Nature Methods 14, 290–296 (2017) 10.1038/nmeth.4169

[57] Sanchez-Garcia, R., Gomez-Blanco, J., Cuervo, A., Carazo, J.M., Sorzano, C.O.S., Vargas, J.: Deepemhancer: a deep learning solution for cryo-em volume post-processing. Communications biology 4(1), 874 (2021)

[58] Cao, Y., Yisimayi, A., Jian, F., Song, W., Xiao, T., Wang, L., Du, S., Wang, J., Li, Q., Chen, X., Yu, Y., Wang, P., Zhang, Z., Liu, P., An, R., Hao, X., Wang, Y., Feng, R., Sun, H., Zhao, L., Zhang, W., Zhao, D., Zheng, J., Yu, L., Li, C., Zhang, N., Wang, R., Niu, X., Yang, S., Song, X., Chai, Y., Hu, Y., Shi, Y., Zheng, L., Li, Z., Gu, Q., Shao, F., Huang, W., Jin, R., Shen, Z., Wang, Y., Wang, X., Xiao, J., Xie, X.S.: Ba.2.12.1, ba.4 and ba.5 escape antibodies elicited by omicron infection. Nature 608(7923), 593–602 (2022) 10.1038/s41586-022-04980-y

[59] Du, S., Cao, Y., Zhu, Q., Yu, P., Qi, F., Wang, G., Du, X., Bao, L., Deng, W., Zhu, H., Liu, J., Nie, J., Zheng, Y., Liang, H., Liu, R., Gong, S., Xu, H., Yisimayi, A., Lv, Q., Wang, B., He, R., Han, Y., Zhao, W., Bai, Y., Qu, Y., Gao, X., Ji, C., Wang, Q., Gao, N., Huang, W., Wang, Y., Xie, X.S., Su, X.-d., Xiao, J., Qin, C.: Structurally resolved sars-cov-2 antibody shows high efficacy in severely infected hamsters and provides a potent cocktail pairing strategy. Cell 183(4), 1013–102313 (2020) 10.1016/j.cell.2020.09.035

[60] Westendorf, K., Žentelis, S., Wang, L., Foster, D., Vaillancourt, P., Wiggin, M., Lovett, E., Lee, R., Hendle, J., Pustilnik, A., Sauder, J.M., Kraft, L., Hwang, Y., Siegel, R.W., Chen, J., Heinz, B.A., Higgs, R.E., Kallewaard, N.L., Jepson, K., Goya, R., Smith, M.A., Collins, D.W., Pellacani, D., Xiang, P., Puyraimond, V., Ricicova, M., Devorkin, L., Pritchard, C., O’Neill, A., Dalal, K., Panwar, P., Dhupar, H., Garces, F.A., Cohen, C.A., Dye, J.M., Huie, K.E., Badger, C.V., Kobasa, D., Audet, J., Freitas, J.J., Hassanali, S., Hughes, I., Munoz, L., Palma, H.C., Ramamurthy, B., Cross, R.W., Geisbert, T.W., Menacherry, V., Lokugamage, K., Borisevich, V., Lanz, I., Anderson, L., Sipahimalani, P., Corbett, K.S., Yang, E.S., Zhang, Y., Shi, W., Zhou, T., Choe, M., Misasi, J., Kwong, P.D., Sullivan, N.J., Graham, B.S., Fernandez, T.L., Hansen, C.L., Falconer, E., Mascola, J.R., Jones, B.E., Barnhart, B.C.: Ly-cov1404 (bebtelovimab) potently neutralizes sars-cov-2 variants. bioRxiv (2022) 10.1101/2021.04.30.442182 https://www.biorxiv.org/content/early/2022/03/24/2021.04.30.442182.full.pdf

[61] Hansen, J., Baum, A., Pascal, K.E., Russo, V., Giordano, S., Wloga, E., Fulton, B.O., Yan, Y., Koon, K., Patel, K., Chung, K.M., Hermann, A., Ullman, E., Cruz, J., Rafique, A., Huang, T., Fairhurst, J., Libertiny, C., Malbec, M., Lee, W.-y., Welsh, R., Farr, G., Pennington, S., Deshpande, D., Cheng, J., Watty, A., Bouffard, P., Babb, R., Levenkova, N., Chen, C., Zhang, B., Hernandez, A.R., Saotome, K., Zhou, Y., Franklin, M., Sivapalasingam, S., Lye, D.C., Weston, S., Logue, J., Haupt, R., Frieman, M., Chen, G., Olson, W., Murphy, A.J., Stahl, N., Yancopoulos, G.D., Kyratsous, C.A.: Studies in humanized mice and convalescent humans yield a sars-cov-2 antibody cocktail. Science 369(6506), 1010–1014 (2020) 10.1126/science.abd0827 https://www.science.org/doi/pdf/10.1126/science.abd0827

[62] Park, Y.-J., Marco, A.D., Starr, T.N., Liu, Z., Pinto, D., Walls, A.C., Zatta, F., Zepeda, S.K., Bowen, J.E., Sprouse, K.R., Joshi, A., Giurdanella, M., Guarino, B., Noack, J., Abdelnabi, R., Foo, S.-Y.C., Rosen, L.E., Lempp, F.A., Benigni, F., Snell, G., Neyts, J., Whelan, S.P.J., Virgin, H.W., Bloom, J.D., Corti, D., Pizzuto, M.S., Veesler, D.: Antibody-mediated broad sarbecovirus neutralization through ace2 molecular mimicry. Science 375(6579), 449–454 (2022) 10.1126/science. abm8143 https://www.science.org/doi/pdf/10.1126/science.abm8143

[63] Huang, M., Wu, L., Zheng, A., Xie, Y., He, Q., Rong, X., Han, P., Du, P., Han, P., Zhang, Z., Zhao, R., Jia, Y., Li, L., Bai, B., Hu, Z., Hu, S., Niu, S., Hu, Y., Liu, H., Liu, B., Cui, K., Li, W., Zhao, X., Liu, K., Qi, J., Wang, Q., Gao, G.F.: Atlas of currently available human neutralizing antibodies against sars-cov-2 and escape by omicron sub-variants ba.1/ba.1.1/ba.2/ba.3. Immunity 55(8), 1501–15143 (2022) 10.1016/j.immuni.2022.06.005

[64] Watson, J.L., Juergens, D., Bennett, N.R., Trippe, B.L., Yim, J., Eisenach, H.E., Ahern, W., Borst, A.J., Ragotte, R.J., Milles, L.F., et al.: De novo design of protein structure and function with rfdiffusion. Nature 620(7976), 1089–1100 (2023)

